# Axons morphometry in the human spinal cord

**DOI:** 10.1101/282434

**Authors:** Tanguy Duval, Ariane Saliani, Harris Nami, Antonio Nanci, Nikola Stikov, Hugues Leblond, Julien Cohen-Adad

## Abstract

Due to the technical challenges of large-scale microscopy and analysis, to date only limited knowledge has been made available about axon morphometry (diameter, shape, myelin thickness, density), thereby limiting our understanding of neuronal microstructure and slowing down research on neurodegenerative pathologies. This study addresses this knowledge gap by establishing a state-of-the-art acquisition and analysis framework for mapping axon morphometry, and providing the first comprehensive mapping of axon morphometry in the human spinal cord.

We dissected, fixed and stained a human spinal cord with osmium, and used a scanning electron microscope to image the entirety of 24 axial slices, covering C1 to L5 spinal levels. An automatic method based on deep learning was then used to segment each axon and myelin sheath which, producing maps of axon morphometry. These maps were then registered to a standard spinal cord magnetic resonance imaging (MRI) template.

Between 500,000 (lumbar) and 1 million (cervical) myelinated axons were segmented at each level of this human spinal cord. Morphometric features show a large disparity between tracts, but remarkable right-left symmetry. Results confirm the modality-based organization of the dorsal column in the human, as been observed in the rat. The generated axon morphometry template is publicly available at https://osf.io/8k7jr/ and could be used as a reference for quantitative MRI studies. The proposed framework for axon morphometry mapping could be extended to other parts of the central or peripheral nervous system.

## Introduction

The so-called “white matter” in the central nervous system is composed of axons, a kind of “electric cable” that transmits neural impulses. Surrounded by a myelin sheath that enables faster conduction at higher firing rates, these fibers are grouped into bundles that connect different regions of the brain with the peripheral nervous system. Myelinated and unmyelinated fibers together occupy about 60% of the white matter volume ^1,2^, and the rest is occupied mostly by glial cells and blood vessels. The morphometry of these fibers is remarkably heterogeneous: while white matter can be thought of as a network of identical cables transmitting action potentials, it appears that the size of these fibers can in fact vary from between 0.1 and 10µm ^3^. It also appears that different white matter pathways have different microstructural characteristics ^4^, but to date this observation has mostly been qualitative, and there are limited data that clearly describe and quantify these differences, and how much these fibers vary across human populations in the brain ^5^ and spinal cord ^6,7^.

Traumatic injuries or neurodegenerative diseases such as multiple sclerosis can damage the axons, potentially leading to chronic pain and functional deficits such as paraplegia. The limited knowledge that we have about white matter microstructure is problematic for understanding what happens at the micro- and macroscopic levels under pathological conditions such as multiple sclerosis. Also, non-invasive microstructure imaging methods, used for the diagnosis and prognosis of these pathologies ^8–10^, need quantitative information of the microstructure in order to validate their accuracy.

If we look specifically at the spinal cord, recent anatomy books ^4,6,11^ suggest that knowledge about fiber organisation in the white matter is an accumulation of decades of research by neuroanatomists and it is not uncommon to use as a references studies that are over fifty years old ^12–15^. Indeed, mapping and classifying the connection and the morphometry of millions of fibers requires particularly painstaking studies that cannot easily be reproduced. While the entire spinal cord cytoarchitecture has been described qualitatively ^4^, fiber density and size have been measured quantitatively only in specific tracts. Furthermore, because each tract is generally studied by different authors who employ different methodologies (tissue preparation, imaging system), a quantitative inter-tract comparison is difficult. Another major issue is that the methodology of these pioneer studies is questionable : for instance, the staining based on silver impregnation prevents researchers from differentiating axon and glial processes ^16^, leading to an overestimation of the axonal density (axons per mm^2^) by a factor 10 to 20 ^16^. An extensive review of axon morphometry in the spinal cord can be found in Saliani et al. 2017 ^7^.

Recently though, thanks to the increasing computational power that allows for the acquisition and storage of increasingly bigger datasets, we have seen the emergence of large field of view imaging (e.g. full mouse brain) at a single-axon resolution^17–19^, including the development of automatic software to segment these datasets^20–23^.

In this work, we establish an acquisition and analysis framework for mapping axons in the central nervous system and demonstrate its application in an ex vivo human spinal cord. Following post mortem extraction of the cord, 24 axial images were acquired using a scanning electron microscope (SEM) at a resolution of 130-260 nm/px, resulting in images comprised of approximately 10 gigapixels, covering the cervical to lumbar spinal levels. Axons and their myelin sheaths were automatically segmented using a novel deep learning method ^23^ in order to derive quantitative measures of axonal diameter, axonal density and myelin density. Finally, global statistics were performed to study the differences between levels, left *versus* right tracts and ascending *versus* descending (sensory/motor) tracts. Maps of axon morphometry were combined and registered to a spinal cord template to create the first microstructure atlas of the human spinal cord white matter.

## Results

### Whole-slice imaging and overall results

For all cervical levels (except C8), four thoracic and six lumbar parts were imaged in mosaic, stitched and segmented (see supplementary material S1).

A remarkable diversity of microstructure could be observed. For instance, we report a much larger axonal density in the fasciculus proprius (30% at C3, blue boxes in figure 1) than in the gracilis regions (15-20%, green boxes), and the presence of larger axons in the corticospinal tract (3.2µm, red boxes) than in the gracilis tract (2.5µm).

**Figure 1.**
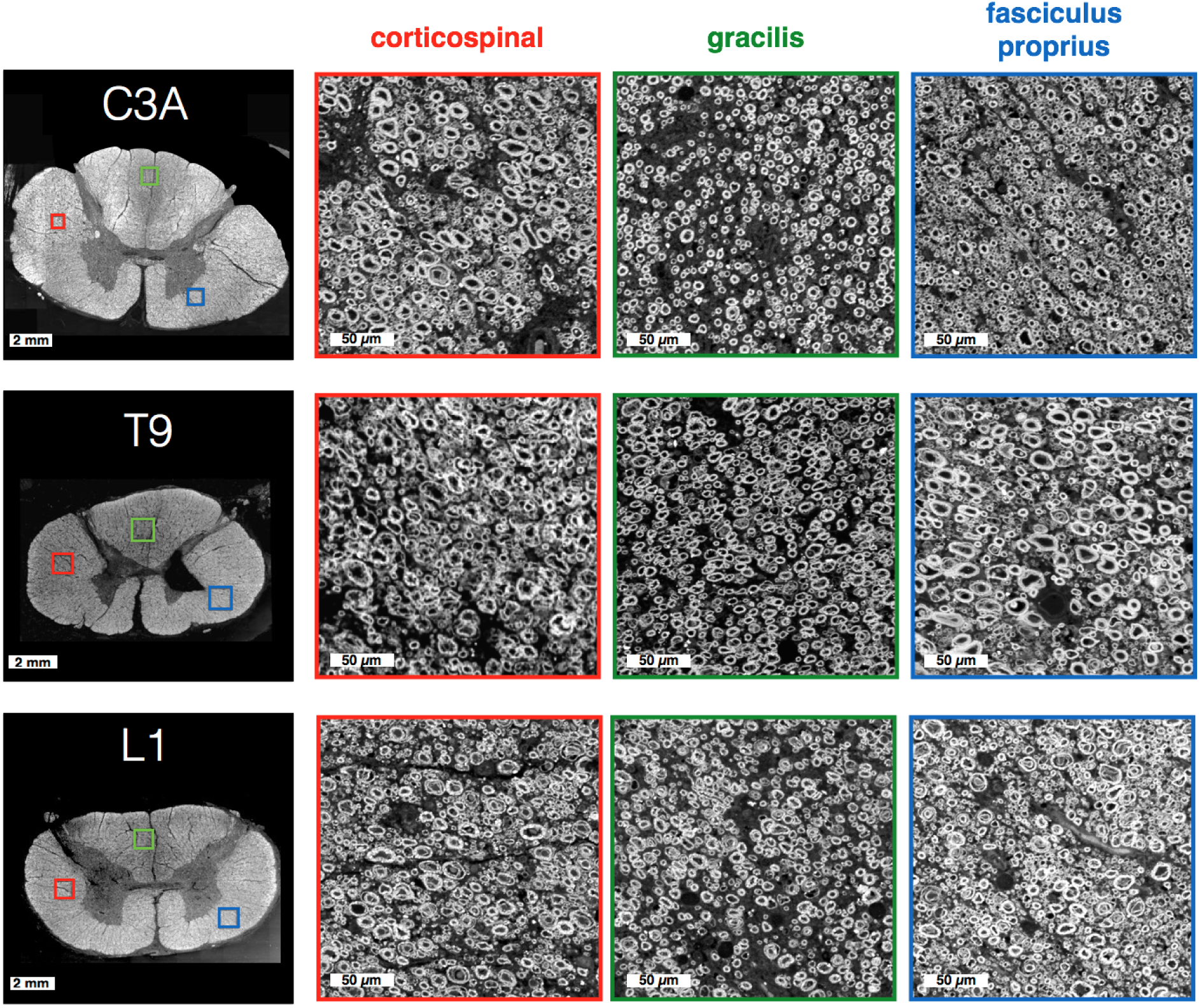
SEM images of the spinal cord over three pathways (color-coded in red, green and blue), at three spinal levels (C3, T9 and L1). Myelinated axons can easily be distinguished on these raw SEM images.

### Axon and myelin segmentation

Image and segmentation quality were first qualitatively assessed in several regions. Myelin sheaths were clearly identifiable with relatively sharp borders (figure 2b), and the large majority of the axons were detected (figure 2c). Using manual segmentation as a ground truth over a small portion of the image (1 x 0.5 mm^2^), the performance of the automatic axon segmentation was evaluated as yielding a sensitivity of 87% and a precision of 78%.

**Figure 2.**
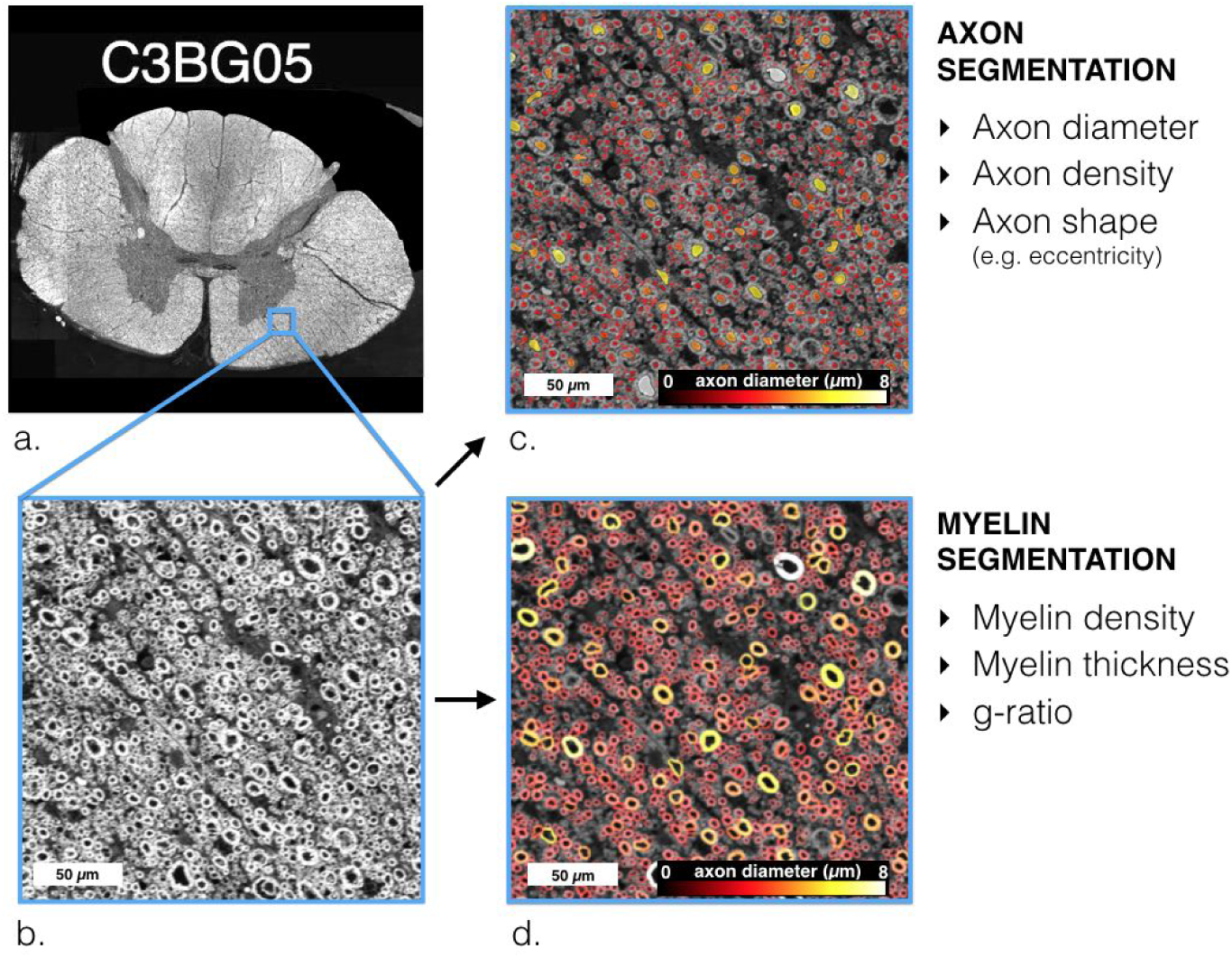
Extraction of microstructural information. a: Large-scale SEM image after stitching all sub-windows, b: zoomed window of 100×100µm^2^, c: segmentation of axons and d: segmentation of myelin. The computed morphometric information is listed on the right panel.

### Microstructural maps

Figure 3 shows the microstructural maps obtained by downsampling the segmented histological images at a resolution of 100×100µm^2^ so as to compute measures of fractional volume (axon and myelin volume fraction) as well as to aggregate measures per unit surface (axon diameter, eccentricity, etc.) The resulting maps were consistent across the different slices (see supplementary material S2) and were remarkably symmetrical (see section “Laterality difference”).

**Figure 3:**
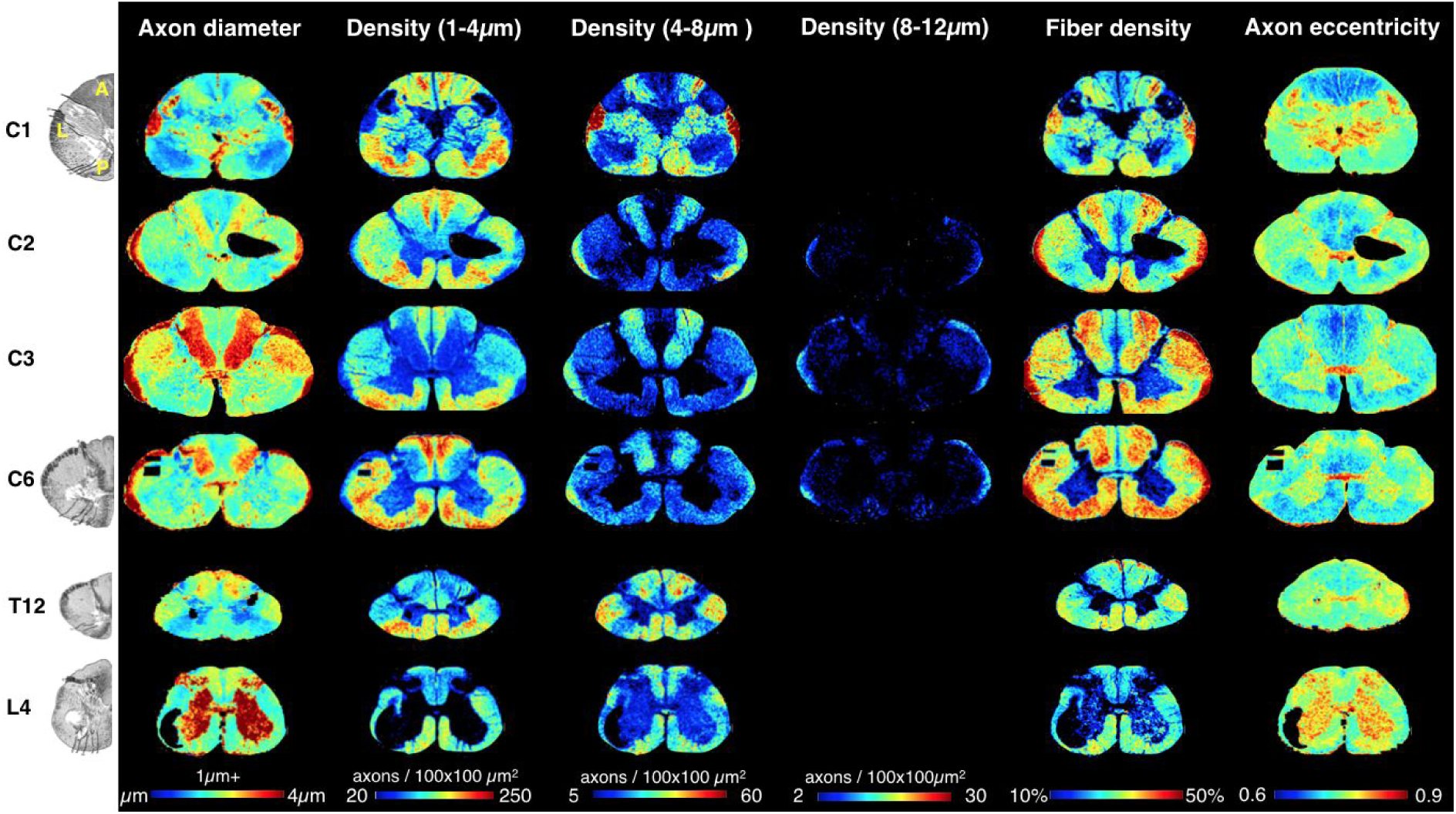
Maps of axon microstructure. Mean axonal diameter (first column), 1-4µm density defined as the number of axons per 100×100 µm^2^ area unit (second column), 4-8µm density (third column), 8-12µm density (fourth column), fiber (axon and myelin) density (fifth column), and mean axonal eccentricity (sixth column) in pixels of 100×100 µm^2^ at different spinal levels (rows). These maps can be compared with the manual drawing of cytoarchitecture extracted from ^4^ (left).

The axon diameter metric was divided into three density maps (1-4µm axons, 4-8µm and 8-12µm, reported as number of axons in a 100×100µm^2^ window). This subdivision clearly reveals the large proprioceptive axons of the cuneatus and spinocerebellar pathways ^24^. Nearly no axons (less than 5 per 100×100µm^2^ window) larger than 8µm were detected, except in the lateral border of the cervical spinal cord (which corresponds approximately to the spinocerebellar pathway).

Axon eccentricity maps represent a combination of two effects: non perfectly perpendicular cutting with respect to the axon orientation (and thus information about the 3rd dimension) and axonal compression. Maps of axonal eccentricity were highly symmetrical, suggesting that the contrast within the spinal cord is mostly driven by genuine microstructural characteristics (i.e. not compression or cutting artefacts). Some tracts (e.g. dorsal columns) present axons that are closer to the shape of a circle, indicating that the axons are running straight along the spinal cord. On the contrary, axons in the corticospinal tract are more oblique.

The gray matter presents a very different microstructure than the white matter in terms of axonal density and orientation. Because the segmentation software and the correction framework was trained on white matter regions, the accuracy of the measures is compromised. However, a comparison against manual segmentation shows overall satisfactory results over the gray matter as well. See Supplementary material S8 for more details.

### Atlas-based analysis

Axon morphometrics were then extracted for each tract using an atlas registration method (figure 9, online methods). We investigated the variation of axon morphometry with respect to the spinal cord pathway, spinal level and laterality.

#### Inter-tract variability

In the white matter section of level C5B (figure 4), the mean axon diameter ranged from 2.5µm in the gracilis tract to 3.4µm in the spinocerebellar tract. Axonal density varied between 40% in the gracilis tract and 50% in the spino-olivary tract. Although the distribution of microstructure can be very broad in a few tracts (e.g. spinocerebellar) due to the presence of outliers (corresponding to the pixels with partial voluming with the gray matter or the background), the 95% confidence intervals are very narrow and reveal significant microstructural differences between tracts.

**Figure 4:**
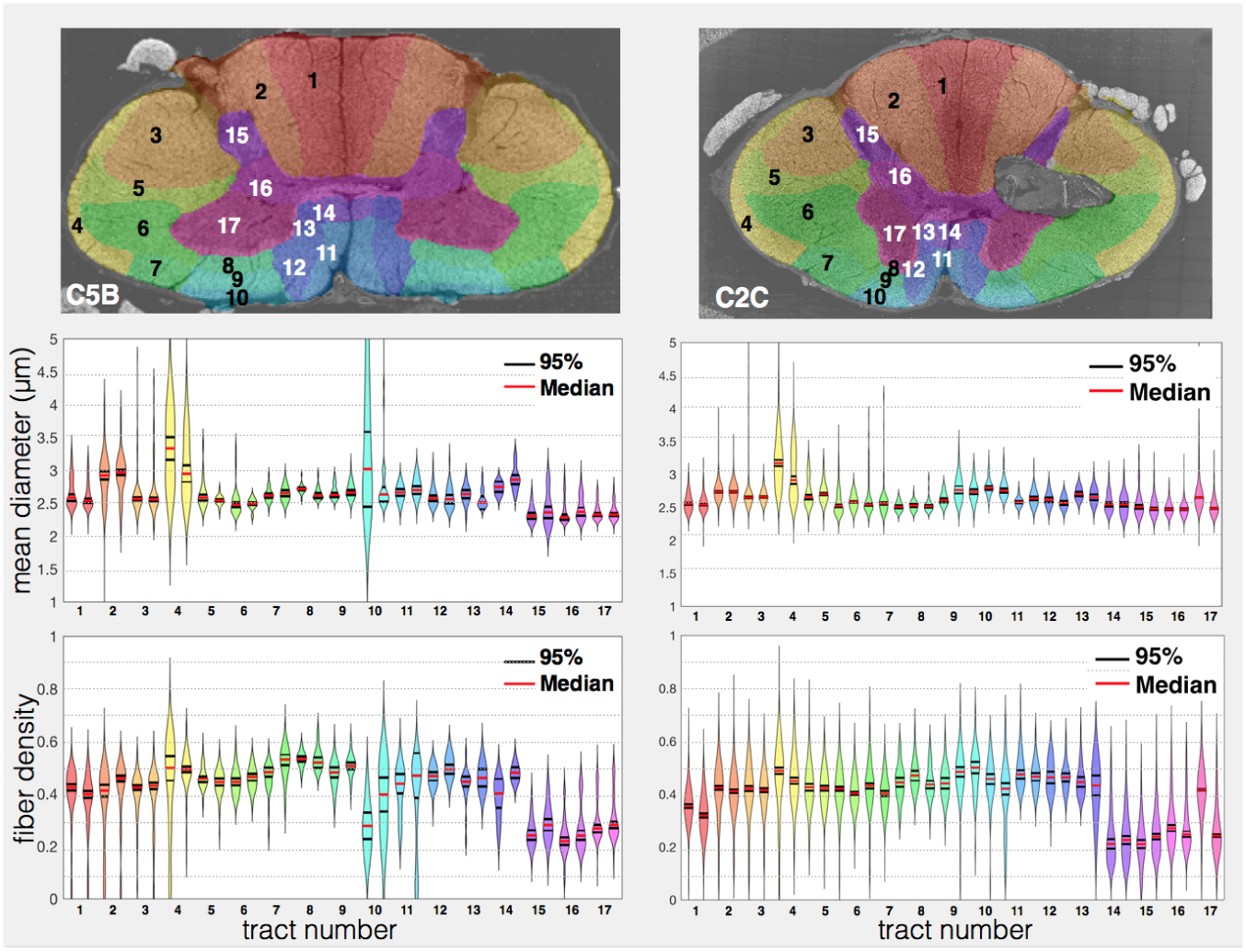
Atlas-based analysis. The spinal cord white matter atlas, built from Gray’s Anatomy ^11^ and available in the Spinal Cord Toolbox ^25^ was registered to the histological slices in order to extract axonal morphometry in the following spinal cord pathways (listed in abscissa as “tract number”): 1: gracilis, 2: cuneatus, 3: lateral corticospinal, 4: spinocerebellar, 5: rubrospinal, 6: spinal lemniscus, 7: spino-olivary, 8: ventrolateral reticulospinal, 9: lateral vestibulospinal, 10: ventral reticulospinal, 11: ventral corticospinal, 12: tectospinal, 13: medial reticulospinal, 14: medial longitudinal fasciculus, 15: Gray Matter dorsal horn, 16: Gray matter intermediate zone, 17: Gray matter ventral horn. Distribution of the mean axonal diameter (middle row) and fiber density (bottom row) within each pathway is shown. The median (red lines) and 95% confidence interval (black lines) show that each tract presents very singular microstructures. Most tracts exhibit a fairly homogeneous microstructure as shown by the narrow distributions.

#### Effect of spinal level

Slice position was defined in the spinal cord template coordinates ^26^. Morphometric properties were averaged in the white matter (all tracts of the registered atlas) and plotted as a function of the slice position (see figure 5). Values were consistent across slices and a smoothing spline was fitted to highlight the trends. Note that T4 was found to be an outlier because of poor focus during the SEM acquisition and was not used in these results (see discussion).

**Figure 5:**
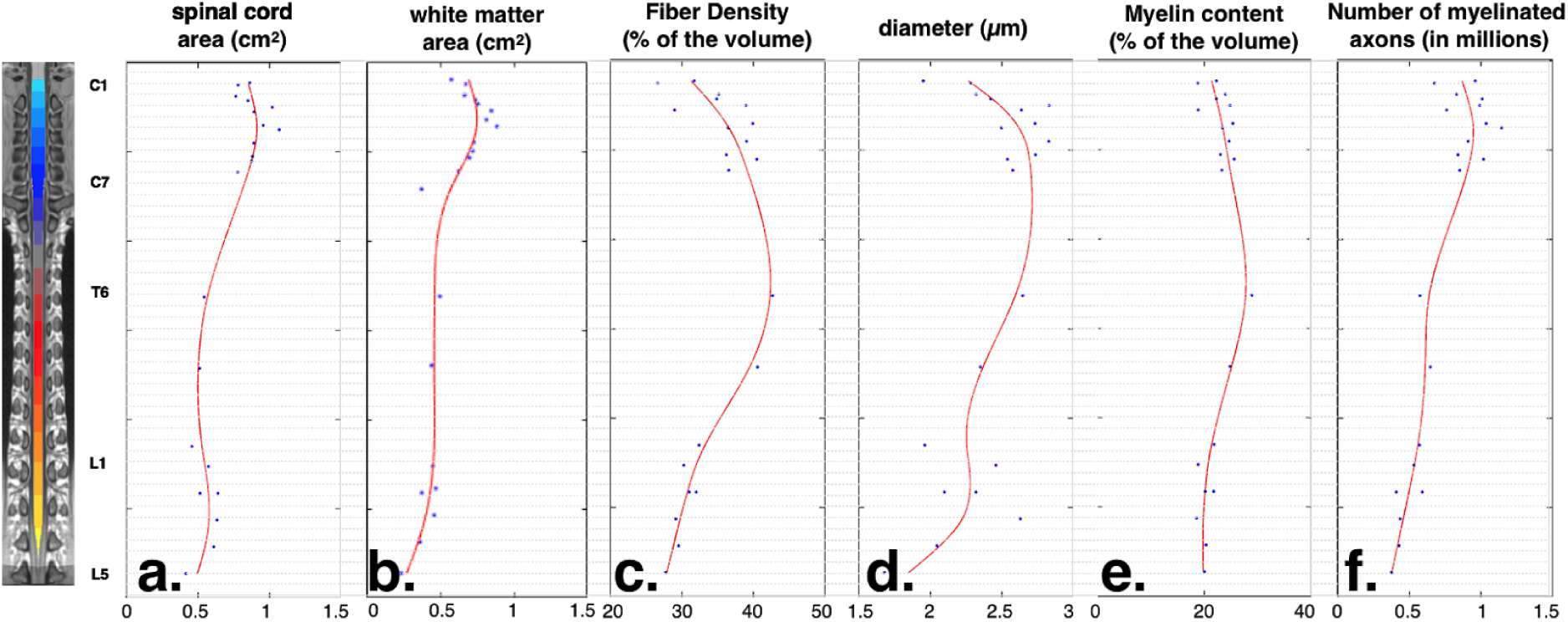
Evolution of the axonal morphometry along the spinal cord. Morphometric properties were averaged in the white matter for each sample (blue dots), and a smoothing spline (red line) was fitted for visualization.

The number of axons increases monotonically towards the rostral direction until about C3, which corresponds to the arrival of afferent fibers at each spinal level. This increase in the number of fibers (Figure 5, f.) correlates with the enlargement (a.) of the spinal cord (r=0.9, p=10^-7^) more than it correlates with the fiber density (c.) (r=0.5, p=10^-2^). The spinal cord cross-sectional area (a.) also correlates significantly with the average diameter (d.) of the fibers (r=0.7, p=10^-4^). The myelin content (i.e. myelin volume fraction) (e.) along the spinal cord is relatively constant, with a difference of only 3% between the lumbar (average of 20%) and cervical parts (23%). Axonal diameter, fiber density and myelin content present maximal values between low cervical levels and mid thoracic level.

#### Laterality difference

The spinal cord microstructure was found to be remarkably symmetrical. When comparing the left and right tracts on a Bland-Altman plot (see supplementary material S9), we found an average difference between left and right tracts of 0.01µm for axon diameter, 1% for axon density and 0.4% for myelin content. This result suggests that handedness has very little impact on axonal morphometry, which needs to be confirmed in a larger sample study.

### Template

The microstructural maps were registered to the PAM50 spinal cord template developed by De Leener et al. ^26^ in order to generate the first MRI-compatible open-access microstructural template of the human cervical spinal cord (see figure 6). The combination of regularized SyN transformations and multi-resolution iterative registrations (from 8 to 2 downsampling factors, mesh size of 12) resulted in fairly smooth transitions between adjacent slices. The template covers C1 to C8 spinal levels. A large transition of microstructure and gray matter shape, as well as a lower sampling rate, led to unsatisfying results for levels below C8, hence they were not included in the template.

**Figure 6.**
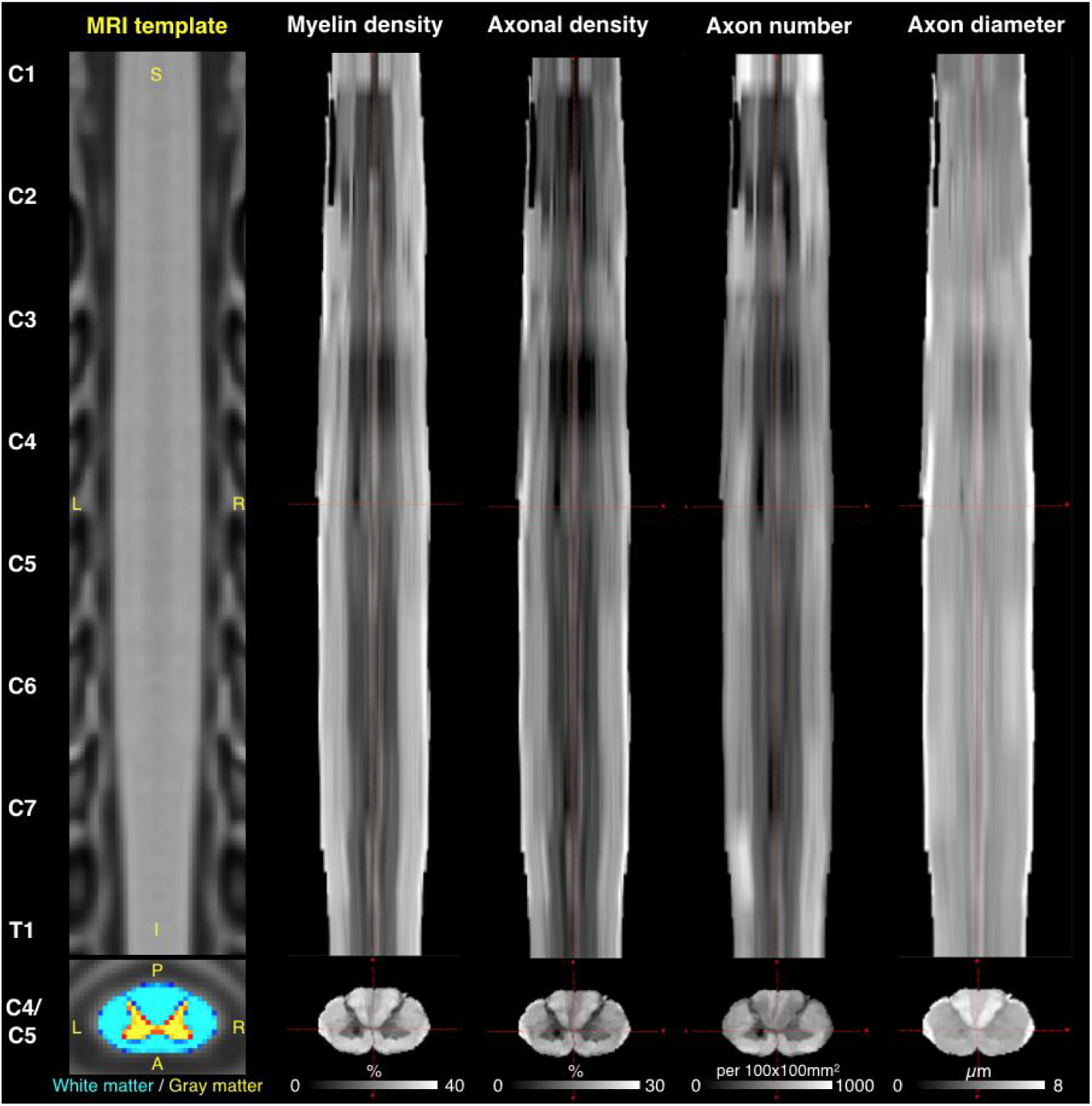
Microstructural template of the spinal cord. Coronal (top) and axial (bottom) view of the generated template. Histological slices were registered at the corresponding vertebral levels of the PAM50 MRI template (left), and then interpolated between slices using non-linear deformations. The resulting template shows fairly smooth transitions between slices. Note that “axon number” refers to myelinated axons only. The white/gray matter overlaid on the axial view of the MRI template is a probabilistic representation of the spinal cord internal structure ^25^ and shows similar morphometry when compared to the histology-based atlases.

## Discussion

In this work we combined high resolution whole-slice histology and automatic axon and myelin segmentation software to produce the first white matter morphometric maps of the entire human spinal cord. While the spinal cord is highly symmetric, different spinal cord pathways exhibit distinct microstructural features.

### Axon morphometry in the spinal cord

A question that immediately arises is why we observe such substantial variations of microstructure between the different spinal cord regions? For example, axon diameters have been observed to vary between 0.1 to 10µm ^3^. Using bioelectrical models and experimental setups, it can be shown that, in myelinated axons, the speed of propagation and the firing rate are proportional to the axon diameter (d) ^27^; while the energy consumption and the fiber volume are proportional to d^2 3^. The presence of large axons in the CNS comes at the expense of energy and space efficacy, and must be compensated by significant benefits. The need for faster propagation in the peripheral nervous system can be appreciated in terms of the advantage gained by faster reflexes (e.g. squid giant axon), or to ensure the synchronization of the neural impulses that come from organs located at various distances from the brain. However, the need for faster propagation cannot generally explain the presence of large myelinated axons in the central nervous system (CNS) for several reasons: (i) axon diameter does not scale with animal size ^28,29^, (ii) there is no correlation between axonal length and axonal diameter, notably in the optic nerve ^2^, (iii) axon diameter in the spinal cord increases towards the rostral direction (see figure 5). Therefore, the second hypothesis, which considers the firing rate as the main reason for larger axons in the CNS, appears to be more likely ^3^.

Electromyography studies of the peripheral axons connected to the spinal cord show a variety of firing frequencies and behaviors (brief bursts or continuous fire) ^30,31^, and thus yields very diverse information. A possible interpretation is that the information that requires a high bandwidth is transmitted “unfiltered” through the largest axons. Conversely, if the same amount of information (same information-rate) is encoded at a lower bandwidth, this information can be transmitted through multiple small axons for more efficacy ^3^. Based on this hypothesis, the microstructural maps (figure 3), and notably the axon diameter maps, reveal a functional organization. This idea was notably suggested by Niu et al. ^24^, where authors compared the proprioceptive and mechanoreceptive axons of the dorsal column using fluorescence labeling ^24^. They showed that these two types of afferent sensory axons possess a distinct caliber, resulting in the contrast between the gracilis and the cuneatus (see axon diameter maps of figure 3), which is classically interpreted as the result of the “somatotopic organization” of the dorsal column.

In the same study ^24^, authors suggest that the large proprioceptive axons enter the dorsal column through the dorsal roots as low as sacral levels, and terminate about 5 segments upward, whereas smaller mechanoreceptive axons enter the dorsal column at higher levels and terminate directly in the medulla. This model would explain the presence of numerous large axons (4-8 µm) in the dorsal column at lumbar and low thoracic level (figure 3), and their reduction in number as we go rostrally. More research is necessary to confirm this model of the medial lemniscus pathway.

### Challenges of tissue preparation

#### Formaldehyde concentration

The first challenge in all histological studies on post-mortem tissue is the preservation of the tissue and, in particular, the integrity of the myelin. In this work, we used a relatively standard fixation procedure (post-fixation via immersion using a mixture of glutaraldehyde and paraformaldehyde). Various concentrations of glutaraldehyde (0.5-4%) can be found in the literature ^32–34^. Due to the uncertainty on the impact of this concentration on axonal morphometry, we decided to use three concentrations (0, 0.5 and 2%). No particular differences were found between the three fixatives on the metrics reported, as shown by the good consistency across slices (see supplementary material S2 and figure 5).

#### Myelin integrity

Myelin integrity was not perfectly preserved, as shown by the abnormally low g-ratio observed (see supplementary material S3). This is quite inevitable when no perfusion fixation can be performed. This observation motivated the report of metrics that are not impacted much by this issue (see discussion about accuracy).

#### Osmium penetration

Concerning the staining procedure, a few slices presented some spots with bad osmium staining (see supplementary material, figure S1, white arrows). Osmium only penetrates a few hundred microns within the tissue, ^19^ making it difficult to ensure a good uniform staining across such large samples. These spots are the result of the presence, in some cases, of air bubbles in the epoxy resin and due to a surface that wasn’t perfectly flat. In order to prevent the polishing to go deeper than a few hundred microns, the surface was exposed by cutting it at various angles. Another strategy would be to improve the osmium penetration by using recently published protocols ^18,19^.

### Challenges of image acquisition

#### Timing

Using the present method, at a resolution of 130nm, it took about 8 hours to image each slice. This painstaking procedure would make it difficult to scan multiple spinal cords, and hence imaging needs to be accelerated. Multi-beam solutions ^35^ are encouraging strategies for reducing scanning time without compromising image resolution.

#### Focus

The images were obtained in low magnification mode. This choice was motivated by the large field of view and the large depth of field of this mode. Still, two slices (T4 and C2A) suffered from poor focus, which had to be discarded from the analysis. We experienced a loss of focus when the SEM vacuum level was slightly altered. We therefore used liquid nitrogen and refilled [refilled what??] during the scan in order to maintain the stability of the vacuum.

#### Resolution limit

In this study, in order to increase the backscattered signal of the electrons as well as to increase the speed of acquisition, we used a relatively high voltage (i.e. 15 keV). This choice resulted in a reduction of the structural details (due to higher penetration, and thus larger electron interaction volume). We estimated the effective spatial resolution by computing the intensity profile of a few large axons using ImageJ. Using the Rayleigh criterion, the spatial resolution enabled us to distinguish only myelinated axons larger than 1µm internal diameter (i.e. about 8 pixels for the 129nm resolution). As a result, we can observe in figure S5 of supplementary material (manual segmentation), that only axons larger than about 1µm were distinguishable in our images (note the sharp drop in the axon diameter distribution). Unfortunately, the literature does not agree on the presence of numerous myelinated axons smaller than 1µm in the human spinal cord ^7^. A sharp drop in diameter distribution has been reported in the macaque corticospinal tract at about 0.5 µm using transmission electron microscopy ^33^, but the whole distribution is shifted toward smaller axons in the macaque with a peak at 1µm.

Note that optical microscopy was also an option, but getting very thin sections (∼5µm) of the entire human spinal cord without any damage proved to be too challenging. The scanning electron microscopy can still be improved by reducing the voltage and by using the high magnification mode, but that would be at the expense of scanning time, contrast to noise ratio, or shorter depth of field.

#### 2D imaging

In this work, 2D images of the spinal cord were obtained, based on the assumption of relatively straight axons running along the spinal cord (axons orientation is dispersed by 28° in the spinal cord) ^36^. This assumption, however, would not hold in brain regions with a large orientation dispersion; rather 3D scanning electron microscopy ^37,38^ or 3D optical imaging ^21,39^ could be considered in these cases. Note that 3D imaging will also make the dataset a hundredfold larger, thereby rendering acquisition and processing time extremely lengthy.

The robustness of our measurements with regard to the orientation dispersion of the axons has been studied in supplementary material S7. The impact is limited (<10% for diameter measurement) for moderate dispersion of axon orientation (<35%) such as in the spinal cord.

Interestingly, the shape of the axons (i.e. elliptical shapes) provides some information about the third dimension (direction of the fibers). This interpretation could be questioned because axonal compression is a confounding factor that cannot be separated easily. However, maps of axonal eccentricity were highly symmetrical, suggesting that the contrast within the spinal cord reveals microstructural information (as opposed to compression, shearing or cutting artefacts that would result in local or asymmetric contrasts). Note that the axon eccentricity measured in some regions (up to 0.7) correspond to excessively high angles (45°), which shows that axons are not perfectly cylindrical.

### Other challenges

#### Staining

Finally, it should be mentioned that the current protocol is blind to unmyelinated axons and glial cells due to the use of osmium, which only stains for myelin. These structures can be seen at a lower voltage and advanced osmium staining method ^19^, but would result in a lower signal to noise ratio and more complex images, thereby requiring the development of more advanced segmentation softwares. However, the proportion of unmyelinated axons in the spinal cord white matter is less than 1% (at least in the corticospinal tract) according to Firmin et al. ^33^ and Wada et al. ^16^ (in the macaque and human respectively).

#### Inter-subject variability

This study is based on a single spinal cord extracted from a 71 year old woman. The microstructure is known to change as a function of age ^6,40^ and sex ^41^: for instance, lower axonal density has been reported in elderly people ^6^. The framework needs to be applied in other subjects in order to quantify the variation across age and subjects. This is feasible thanks to the large automation of the acquisition and processing.

### Accuracy of the measurement

As summarized in ^7^, axon morphometry measured with histology vary greatly between studies. For instance, the axonal count may vary between 8,000 and 100,000 axons/mm^2^ in the pyramidal tract, or the proportion of axons smaller than 1µm may vary between 30% to 90%. These strong variations result from (i) staining issues, that can cause confounds (e.g. between axons and glial cells) ^16^ or render poorly stained axons hardly visible ^19^, (ii) the resolution limit of the imaging system that make the detection of the smallest axons impossible ^33^ and introduce a constant bias due to the point spread function, and (iii) tissue degradation (e.g. unwarped myelin). Additionally, segmentation performance (sensitivity, specificity, accuracy) and ambiguities in the methods (myelinated-only or all fibers?) make the comparison very hard.

Values reported in this study concern myelinated-only axons that are larger than 1µm (see resolution limit). Segmentation bias was also limited by using an automatic segmentation method combined with a correction framework detailed in supplementary material S2.

#### Axonal count and fiber density

We found between 10,000 and 20,000 axons/mm^2^ in the corticospinal tract (i.e. pyramidal tract), depending on the vertebral level. We thus confirm the observation of ^16^: pioneer studies such as ^29^ overestimated by about 10 times (100,000 axons/mm^2^ was reported) the number of axons due to staining issues. Note that Wada et al. ^16^ reported a value relatively close to our results: about 9,000 axons/mm^2^.

#### Axon diameter

Axon diameter accuracy can be impacted by the over-segmentation of the myelin, the axonal compression, and the orientation of the axons.

The over-segmentation of the myelin resulted in a constant underestimation of the axonal diameter by a fraction of a micron (see discussion about “resolution limit”).

The axon diameter overestimation for oblique axons was formulated in supplementary material S7. For the majority of the axons (angle <40°), the overestimation was limited to about 10%. Also, AxonSeg filters highly oblique axons (70° maximum) based on their minor to major axis ratio.

#### Myelin content and fiber density

The myelin content (i.e. myelin volume fraction) in the white matter (20-30%, see template) was close to what has been measured by other groups using transmission electron microscopy in other brain regions (25-30%) ^1,2,42^. Note that, contrary to transmission electron microscopy studies, we segment the entire spinal cord and not only small white matter regions, and our reported MVF values can be lower because we take into account the presence of veins and fissures.

As discussed earlier (section “Tissue preparation”), the myelin is not perfectly dense, especially for large axons. Fiber density and myelin content were the most impacted metrics by this limitation. In order to estimate the overestimation of myelin content, we computed the myelin content assuming a constant g-ratio of 0.7 for all axons. With this assumption, the myelin content (MVF) can be related to the axon volume fraction (AVF) as follow : *MV F* = (1/*g*^2^1) × *AV F* = 1.04 × *AV F*. With this approach, we found a linear relationship (data not shown) between our reported MVF value and MVF values obtained assuming a g-ratio of 0.7 : *MV F* _*reported*_ = 0.68 × *MV F*_*g*=0.7_. Likewise, the fiber density (FVF = AVF + MVF) can be computed directly from AVF : *FV F* _*reported*_ = 0.76 × *FV F*_*g*=0.7_. As a result, although the reported values for myelin content and fiber density are meaningful for inter-tract and inter-slice study, readers must be aware of this upward bias. Further work will need to improve myelin integrity or use TEM studies on well preserved axons (with dense myelin) to refine this calibration.

## Conclusion

We present the first microstructural template of the human spinal cord based on histology, registered on the spinal cord MRI template PAM50. To generate this template, we developed a framework that includes whole slice electron microscopy, automatic axon segmentation and coregistration of the slices. The template can be used to revise biophysical models and spinal cord atlases, and validate non-invasive techniques that measure spinal cord microstructure.

## Acknowledgements

We would like to thank Micheline Fortin and the other employees of the histology department of the Institute for research in immunology and cancer (University of Montreal, QC, Canada), Diane Gingras from the electron microscopy department of the University of Montreal, as well as Irène Londono, Monica Nelea and Anik Chevrier. Their help and advice for the preparation of the samples greatly contributed to the success of this work. Finally, we would like to thanks the members of the Center for Characterization and Microscopy of Materials (CM)^2^, notably Nicole MacDonald, Philippe Plamondon and Jean-Philippe Masse, for their feedback on the polishing and scanning of the spinal cords using the electron microscope.

## Online methods

### Tissue preparation

#### Ethics

The spinal cord was extracted from the fresh cadaver of a 71 year old woman cadaver (1.60m tall, weighing 55Kg), donated to the anatomy laboratory at the University du Québec at Trois-Rivières by informed consent. All procedures were approved by the local Ethics Committee (SCELERA-15-03-pr01).

#### Dissection

Two hours after death, the spinal cord was dissected and cut in 5-10mm thick transverse sections (see figure 7). The rostro-caudal position was estimated by counting the nerve roots (8 cervical, 12 thoracic and 5 lumbar). The right side of the spine was marked using strings attached to the nerve roots.

**Figure 7.**
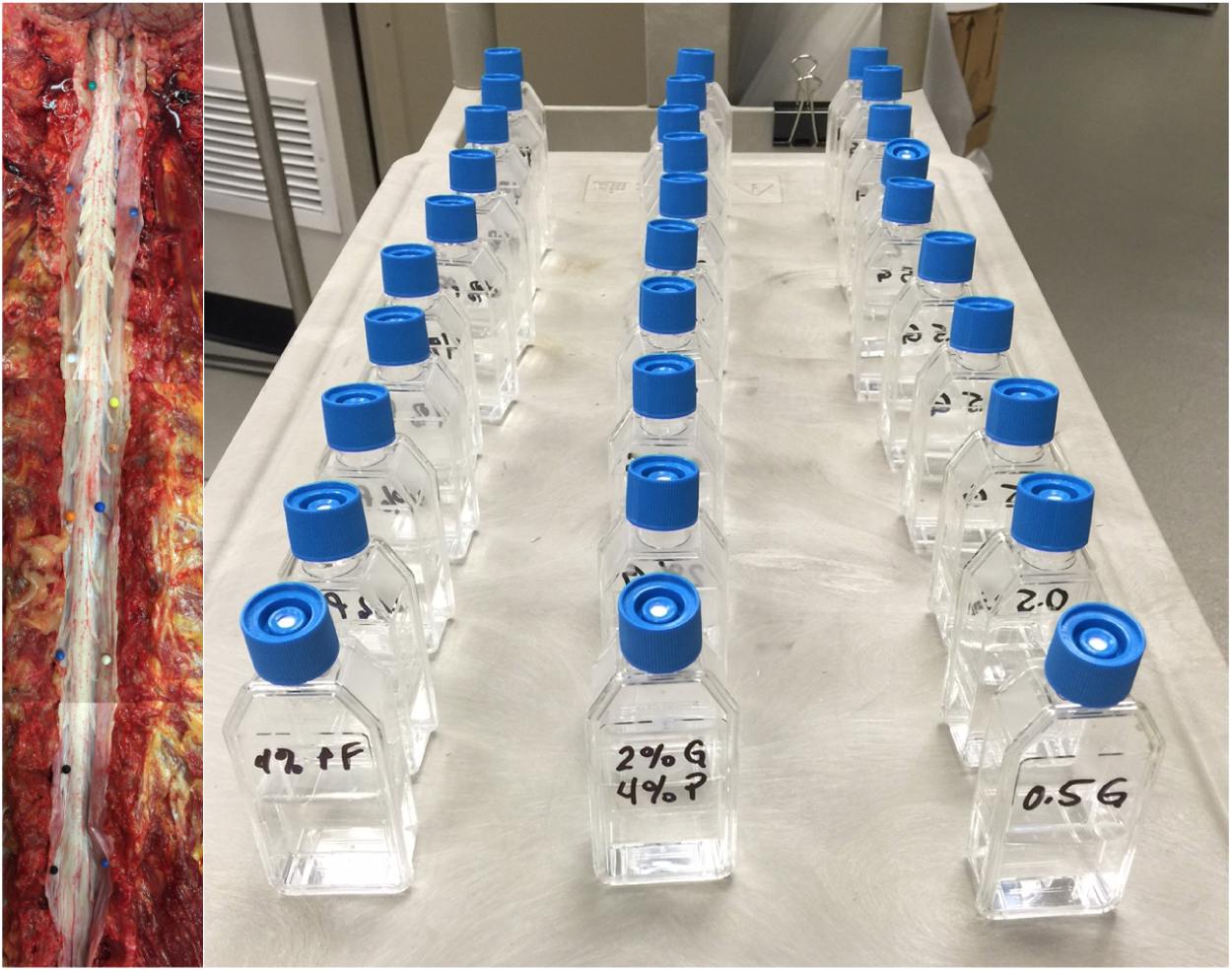
Spinal cord dissection. The spinal cord was first entirely exposed (Left) and 24 one-centimeter thick slices of spinal cord were extracted and post-fixed (right).

#### Fixation

Just after dissection, samples were immersed in separate 50 mL vials containing a solution of 4% paraformaldehyde and 0-2% glutaraldehyde (see table 1), and stored at 4°C. The buffer was a PBS 1x solution, adjusted for a pH of 7.4 with HCl. Different glutaraldehyde concentrations were used in order to assess the bias to tissue shrinkage and prevent the report of wrong conclusions (see discussion). After a week, samples were transferred in PBS 1x to prevent over-fixation. Table 1 lists the extracted spinal cord sections, their spinal level, and the exact glutaraldehyde concentrations.

**Table 1.** List of samples. C: cervical level, T: thoracic level and L: lumbar level. When multiple slices were extracted at the same level, letters A, B and C were used with A the most rostral slice.

| spinal level | PFA | Glutaraldehyde | SEM resolution (nm) |
| --- | --- | --- | --- |
| C1A | 4% | 0% | 260 |
| C1B | 4% | 0% | 260 |
| C2A | 4% | 2% | 130 |
| C2B | 4% | 0% | 260 |
| C2C | 4% | 0.5% | 130 |
| C3A | 4% | 2% | 130 |
| C3B | 4% | 0.5% | 130 |
| C3C | 4% | 0% | 260 |
| C4A | 4% | 2% | 130 |
| C5B | 4% | 0.5% | 130 |
| C5B | 4% | 0.5% | 130 |
| C5B | 4% | 2% | 130 |
| C6A | 4% | 0.5% | 130 |
| C6B | 4% | 2% | 130 |
| C7B | 4% | 0.5% | 130 |
| T12 | 4% | 0% | 260 |
| T4 | 4% | 0% | 130 |
| T9 | 4% | 0.5% | 130 |
| T6 | 4% | 2% | 130 |
| L1 | 4% | 0.5% | 130 |
| L2A | 4% | 2% | 130 |
| L2B | 4% | 0% | 260 |
| L3A | 4% | 0.5% | 130 |
| L4A | 4% | 0% | 260 |
| L5 | 4% | 0% | 260 |

#### Preparation for microscopy

Samples were stained with Osmium 2% during 10h in 10mL vials. A bidirectional rotator was used to prevent deposition of the osmium at the bottom of the vial. Samples were then washed in distilled water, and dehydrated in 10, 25, 50, 75 and 100% acetone baths for 30 minutes each. Acetone was then progressively replaced with 50 and 100% Epon 812 (Mecalab, Canada) for 12 hours each. Final embedding was done at 60°C for 24h. During the final embedding, each axial slice was carefully positioned at the bottom of the mold, and maintained using a plastic grid, in order to have the surface as flat as possible. Once the embedding procedure was finished, a microtome (Reichert-Jung) was used to remove the first layers of resin (15µm thickness) and in order to expose the entire slice of spinal cord (see figure 8). The large tungsten-carbide blade of the microtome could cut the entire block by layers of 15µm. When the cord surface was found to be slightly bent, which is bound to happen with 1 cm large samples, we performed the cutting at various angles. About 50-100µm of tissue was removed before obtaining fully exposed slices of spinal cord. This procedure was necessary because osmium has a penetration depth of only 200µm ^19^. Exposed slices of spinal cord were then polished using a 0.05µm aluminum polishing suspension, and the electrical conduction was ensured using vapor deposition of gold (layer of 600Å).

**Figure 8.**
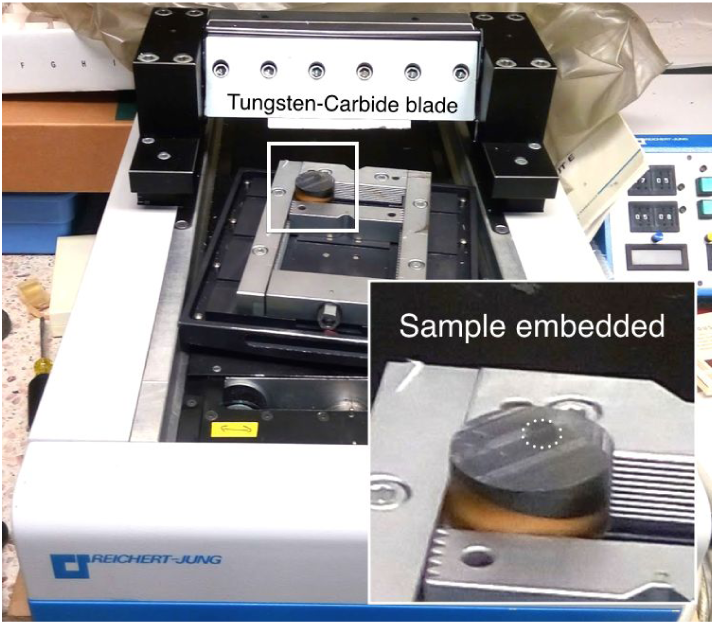
Microtomy. A Tungsten-Carbide blade was used to expose the surface of the spinal cord. The support angle was varied in order to prevent from cutting deeper than 100µm.

### Microscopy

Images were obtained using a scanning electron microscope (SEM) (JEOL JSM7600F) controlled with AZtec 3.2 software (Oxford Instruments, UK). Each sample was carefully positioned on the specimen holder to ensure a parallel surface and good conduction with the specimen holder using carbon tape. The large area mapping solution of AZtec was used to acquire between 150 and 300 mosaics (depending on the spinal level) of 8192×5632 pixels and an area of 1060×729 µm^2^. The following parameters were used for the scanning: low magnification mode, low-angle backscattered electron detector (LABE), aperture of 110 µm, 15 kV, probe-current 140 µA, 110x magnification, 15mm distance, 2µs dwell time. Low magnification mode was selected to increase the field of view and the depth of field. Focus was set in the peripheral nerves to prevent surface degradation, and the contrast/brightness was manually adjusted. If the signal-to-noise ratio was not satisfactory, the scanning distance was reduced to 10mm and the probe current increased to 16.

### Image Processing

#### Stitching

Mosaics were automatically stitched using the *Grid/Collection Stitching Plugin* ^43^ of the image processing program Fiji ^44^. Because the stitching was failing in some regions (usually at the periphery of the spinal cord where the background is present), an outlier detection was implemented assuming a constant shift of the stage (see supplementary material S4).

#### Segmentation

Axons and myelin of each mosaic image were automatically segmented using AxonSeg ^22^ on a computer equipped with a 12 cores Xeon Phi processor. The segmentation of a full slice of spinal cord was obtained in about 12h (using the Matlab parallel computing toolbox). The file containing all segmentation parameters can be found at the following link. For each axon, the following properties were measured:

- Axon equivalent diameter: 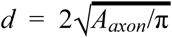 (*A*_*axon*_ is the area of the segmented axon)
- Fiber equivalent diameter: 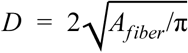 (*A*_*fiber*_ = *A*_*myelin*_ + *A*_*axon*_)
- g-Ratio: 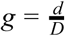
- eccentricity: 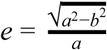 (with a the length of the major axis and b the length of the minor axis of the ellipse that has the same normalized second central moments as the segmented axon ^45, Appendix A^)

In order to prevent over- or under-segmentation, segmentation biases were compensated by comparing manual segmentations (ground truth) and automatic segmentations on a region of 1×0.5mm^2^. Axons smaller than 1µm were considered false positives (see discussion about “resolution limit”), and and were removed from the reported average metrics. See supplementary material S2 for more details.

#### Downsampling

Stitched images were then downsampled at a resolution of 50, 100 and 200µm (resulting in different signal to noise ratio) to produce maps of the *(i)* average axon equivalent diameter, *(ii)* number of axons ranging from 1 to 4µm per pixel, *(iii)* number of axons ranging from 4 to 8µm per pixel, (*iv*) total number of axons per pixel, *(v)* myelin content (i.e. myelin volume fraction), *(vi)* axon volume fraction, and (*vii*) fiber (= myelin plus axon) density (i.e. fiber volume fraction). Note that we extrapolate the volume of myelin or axons from the area by assuming consistency of these areas along the spinal cord axis. Therefore,

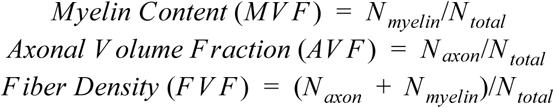

with *N*_*j*_ the number of pixels attributed to class *j* in the downsampling blocks (e.g. for a downsampling from 0.130 to 50µm, *N* _*total*_ = (50/0.13)^2^ = 150, 000 *pixels*).

#### Atlas registration

In order to identify the different spinal cord pathways, we registered the digital version (80×80×500µm^3^) of the Gray’s anatomy ^11^ atlas that is part of the spinal cord toolbox ^25,46^ using elastic deformation to the downsampled 50µm map. For each sample, the corresponding mid-vertebral level (or slightly shifted when multiples slices were extracted at the same level) of the atlas was extracted, and registered in two steps: an initial affine transformation based on manually selected control points (*cpselect* and *fitgeotrans* functions available in the Matlab image processing toolbox), and a diffeomorphic elastic transformation (SyN) estimated on manually drawn masks of spinal cord with two labels for gray and white matter (command *sct_register_multimodal* from the spinal cord toolbox, metric “Mean Squares”) ^46,47^. The elastic transformation (regularized with b-splines, BsplineSyN) was divided into two steps: a first transformation for “smooth” (i.e. global) deformations and a second transformation allowing for more local deformations. Figure 9 shows an example of the registration framework.

**Figure 9:**
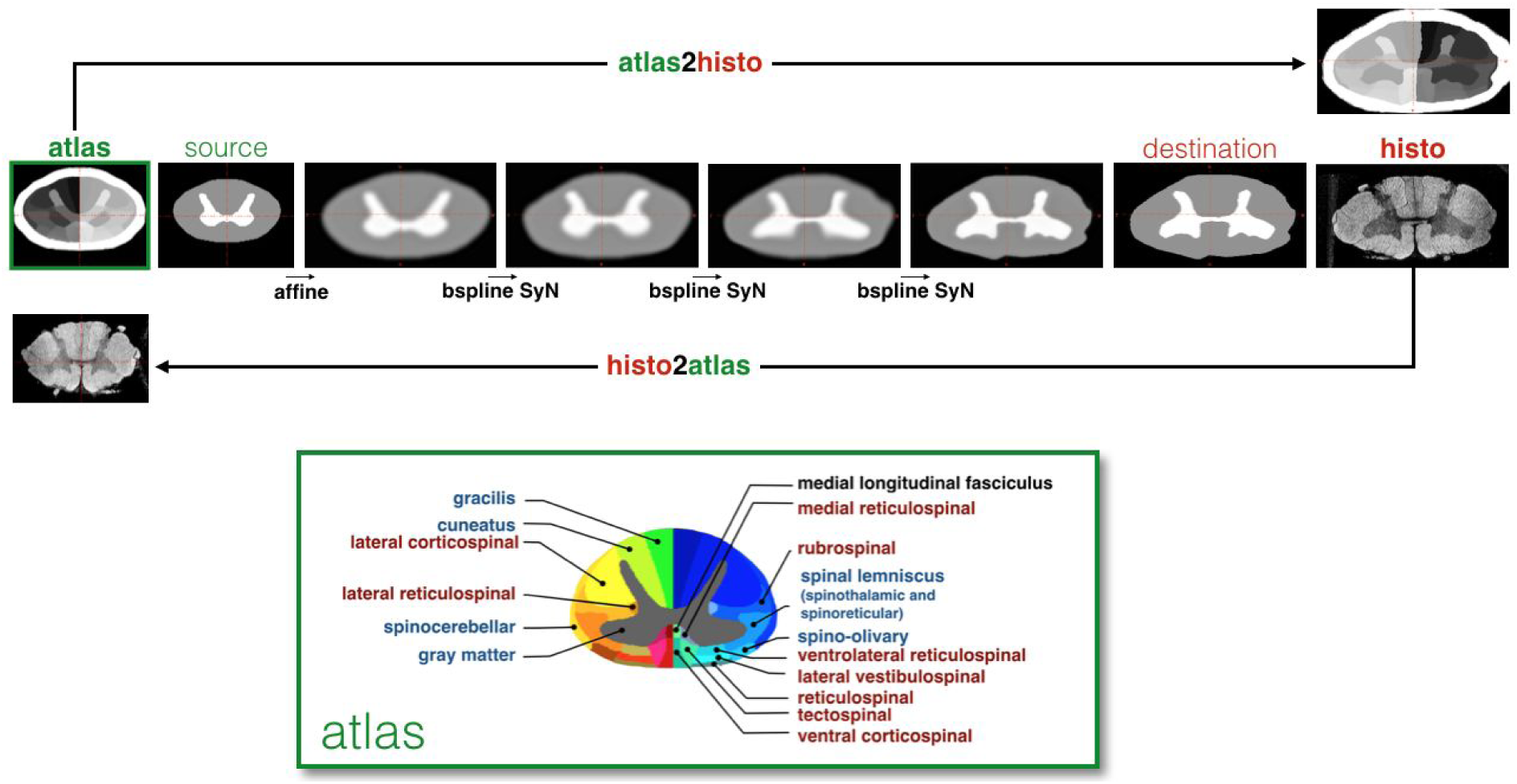
Registration of the atlas (bottom right) to histology (‘histo’). Registration was estimated based on the masks of white and gray matter of the atlas (source) and histology (destination). Multiple steps were used: affine, and multiple non-linear deformations regularized with bsplines (bspline SyN). The inverse transformation (histo2atlas) used only smooth deformation in order to keep the shape of internal structures intact.

#### Template creation

Downsampled maps were registered to the spinal cord PAM50 template (MRI template) ^26^. For this transformation (histology to template space), the local deformation step wasn’t included in order to preserve the shape of the inner structures (e.g. gray matter). Missing levels were filled by registration and interpolation of the closest available slices. The nearest top and bottom slices were first registered to the missing levels using the white matter masks, then the distance-weighted average was used. Due to the small number of slices at thoracic levels, the template was generated for the full cervical part only.

### Statistics

We used the warped spinal cord atlas in the downsampled histology space to measure the axon morphometry in each tract, and then computed correlation between slices by using the Pearson equation. Maximal laterality difference was assessed using the coefficient of reproducibility (1.96 times the standard deviation of the differences), and statistical significance was computed using a paired t-test.

## Supplementary Material

### Whole-slice SEM

**Figure S1.**
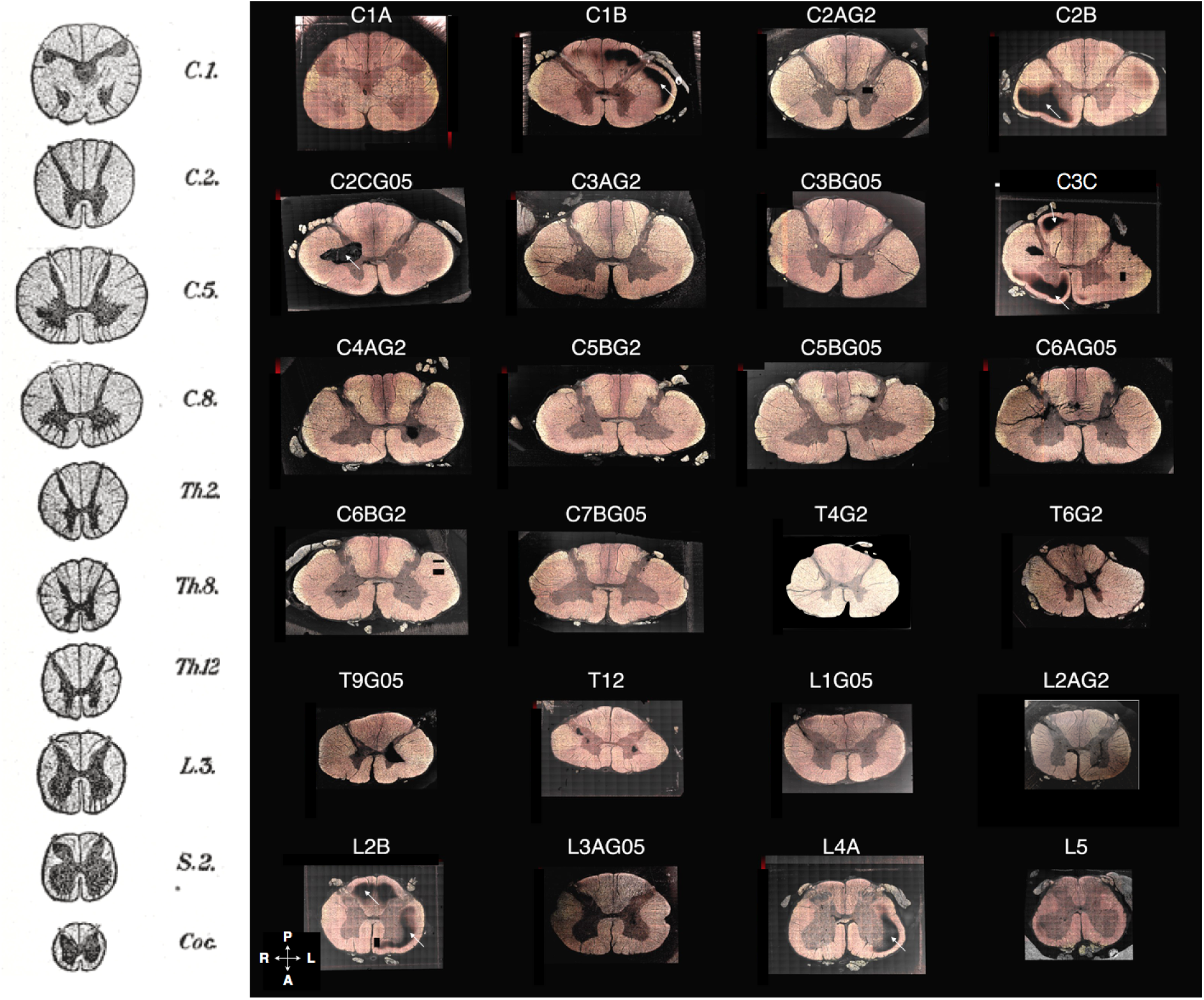
Whole-slice SEM images (stitched), acquired at a resolution up to 129 nm, and segmented using AxonSeg ^22^. The shape of the spinal cord and the grey matter is very similar to the hand drawings from the Gray’s anatomy (Left column) ^11^. P: posterior, A: anterior, L: left, R: right.

### Consistency of the segmentation across slices

Values extracted in the largest tracts (numbered 1 to 6 in figure 4) were consistent between the different cervical levels C2 to C7, whatever the glutaraldehyde concentration used. In these slices, the Pearson’s correlation matrices were 0.86, 0.68, 0.72 on average for axon diameter, myelin volume fraction and axon density respectively. Other slices were less consistent, which is due either to larger microstructural modifications or to strong atlas deformation (e.g. gray matter shape changes rapidly at C1). Figure S2 illustrates the consistency between slices.

**Figure S2.**
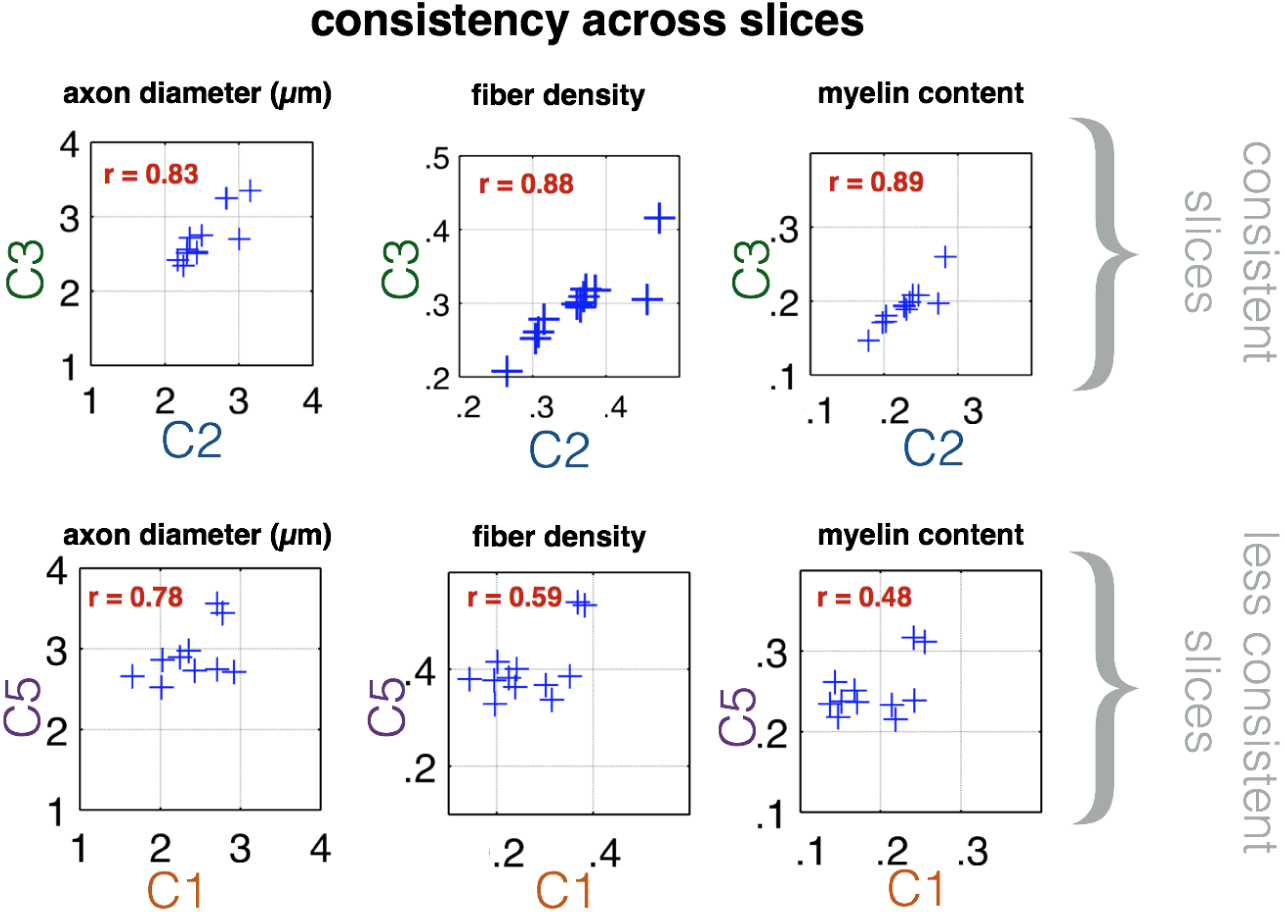
Consistency of the atlas-based analysis across slices (using the largest tracts, numbered 1 to 6 in figure 4). Most cervical slices were strongly consistent (r>0.8) between each other (e.g. C3, 0.5% glutaraldehyde, versus C2, 2% glutaraldehyde, in top row). However, C1 and thoracic levels were less consistent (e.g. C5 versus C1 in bottom row), notably due to particularly strong atlas deformation.

### Axon morphometry distribution within a pixel

Axon morphometry distribution was analysed in small regions (Figure S3). Multiple probability functions were fitted on the axon diameter histogram using allfitdist^1^. The generalized extreme value distribution was found to best describe the histogram with three parameters (shape k, location µ and scaling σ). With only two parameters (location µ and scaling σ), the lognormal distribution was the best. The axon diameter probability density function (pdf) was then weighted by the axon area (π(d/2)^2^) in order to get the fiber volume density function (Figure S3, middle plots). In all the regions studied (spinocerebellar, gracilis and corticospinal tract in slice C5B), the highest fiber volume density was reached at 2.2µm (i.e. this population of fibers occupy a large space).

The g-ratio was found to increase as a function of the diameter, with similar behaviour in the different regions. Interestingly, the g-ratio reveals two clusters of fibers (see zoomed window on bottom right) : small axons with thin myelin sheath (cluster 1), and variable sized axons with a wide range of myelin thickness (cluster 2). Multiple interpretation can be made from these results: (i) Myelin is not perfectly dense for a certain population of axons (cluster 2), and especially the largest ones; (ii) axons in cluster 1 that are larger than 2µm present a normal g-ratio (close to 0.7), suggesting well preserved myelin; (iii) axons smaller than 2 µm with well preserved myelin (cluster 1) present a particularly low g-ratio (down to 0.4), suggesting some systematic overestimation of the myelin thickness due to some blurring (related to the point spread function of the SEM system). Due to these two effects (myelin degradation and blurring), the mean g-ratio values were low (around 0.5), compared to the typical 0.7-0.8 values based on various TEM studies reviewed by.

**Figure S3.**
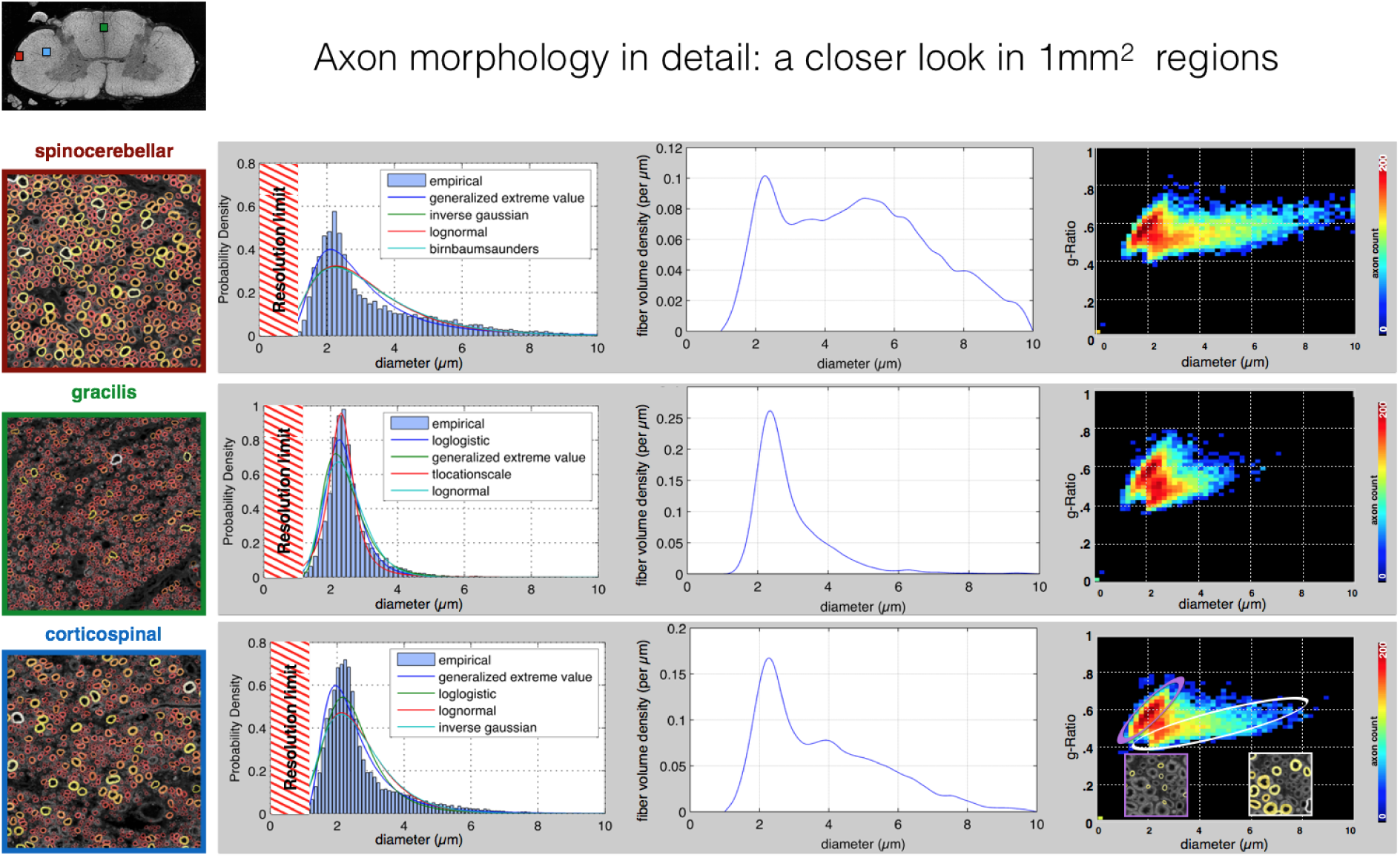
Axon morphometry as a function of axonal diameter in three 1mm^2^ regions (color-coded in red for spinocerebellar, green for gracilis and blue for corticospinal tract) of sample C5B. Left plots: Axon diameter distributions fitted using multiple probability functions (resolution limit was estimated around 1µm, see figure S2). Middle plots:. The fiber volume density as a function of axonal diameter. Right plots.: g-Ratio as a function of axonal diameter.

### Stage displacement correction

Although powerful, the stitching algorithm failed in a few cases, notably for images containing a dark background. This problem was corrected by modeling the stage displacement as two constant shifts:

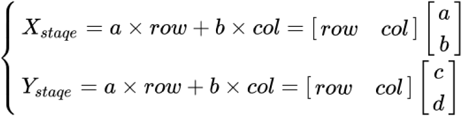

with *row=[1 2 … Nrow]* ^*T*^ and *col=[1 2 … Ncol]*^*T*^ the mosaics index; *X*_*stage*_ and *Y*_*stage*_ the spatial coordinates (in pixels) of the top left corner of each image; a, b, c and d the unknown constants.

Based on the values of *X*_*stage*_ and *Y*_*stage*_ provided by the stitching algorithm, a, b, c and d were estimated (pseudo-inverse solution). Outliers were defined as the coordinates that differ by more than 3 times the median absolute deviation. Figure S4 illustrates the outlier detection algorithm.

**Figure S4.**
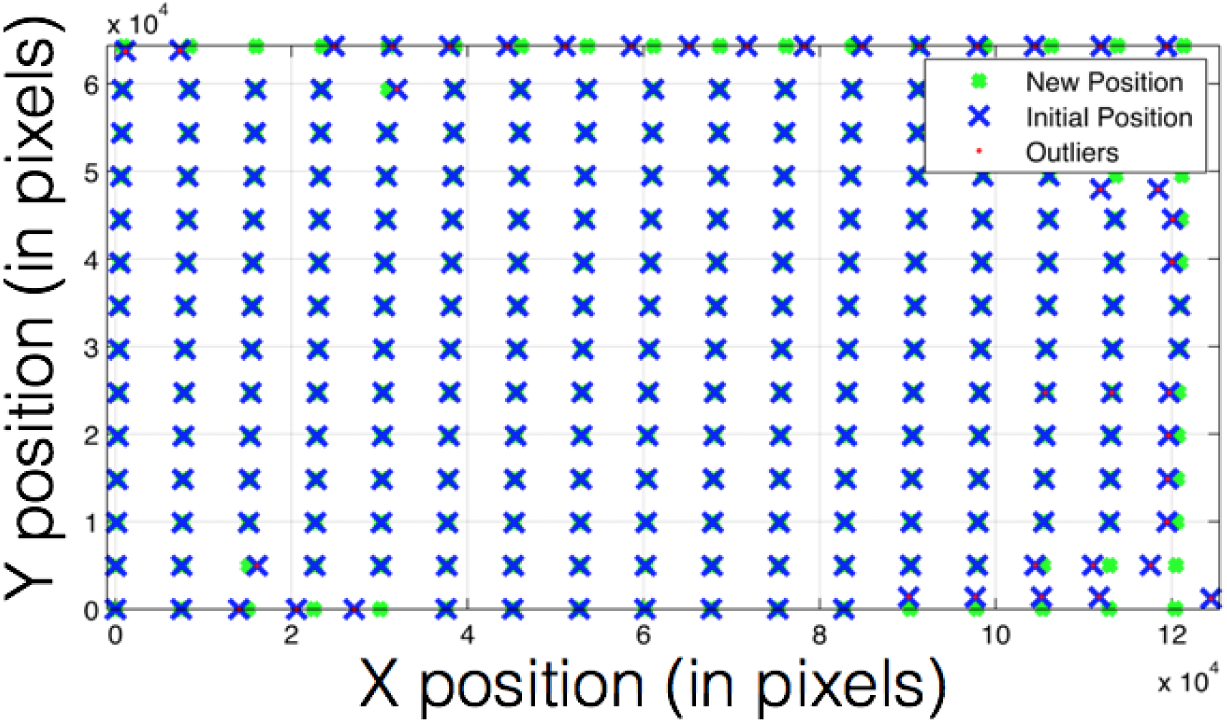
Outliers correction of the stitching algorithm. Mosaics position (blue crosses), estimated using the stitching algorithm, was found to be arranged in a oblique grid (constant bias in X and Y). Based on this assumption, outliers can be detected (red dots) and corrected (green crosses).

### Segmentation calibration framework

Two methods were used in order to correct the axon bias (figure S5). The first method was used to correct the axon density and myelin content by a linear fit. The second method was used to correct the axon diameter and axon number using a histogram correction. The correction was done by doing a training on a manually segmented region (region 1) and comparing this to the automatically segmented image at the same spot. The corrected stats obtained were fitted and then this fit was then applied to a second region on the automatic segmented image (region 2). This was then validated with the groundtruth of this region to ensure that the results were correct. This region field of view was 1×0.5mm^2^ and contained 5133 axons.

**Figure S5.**
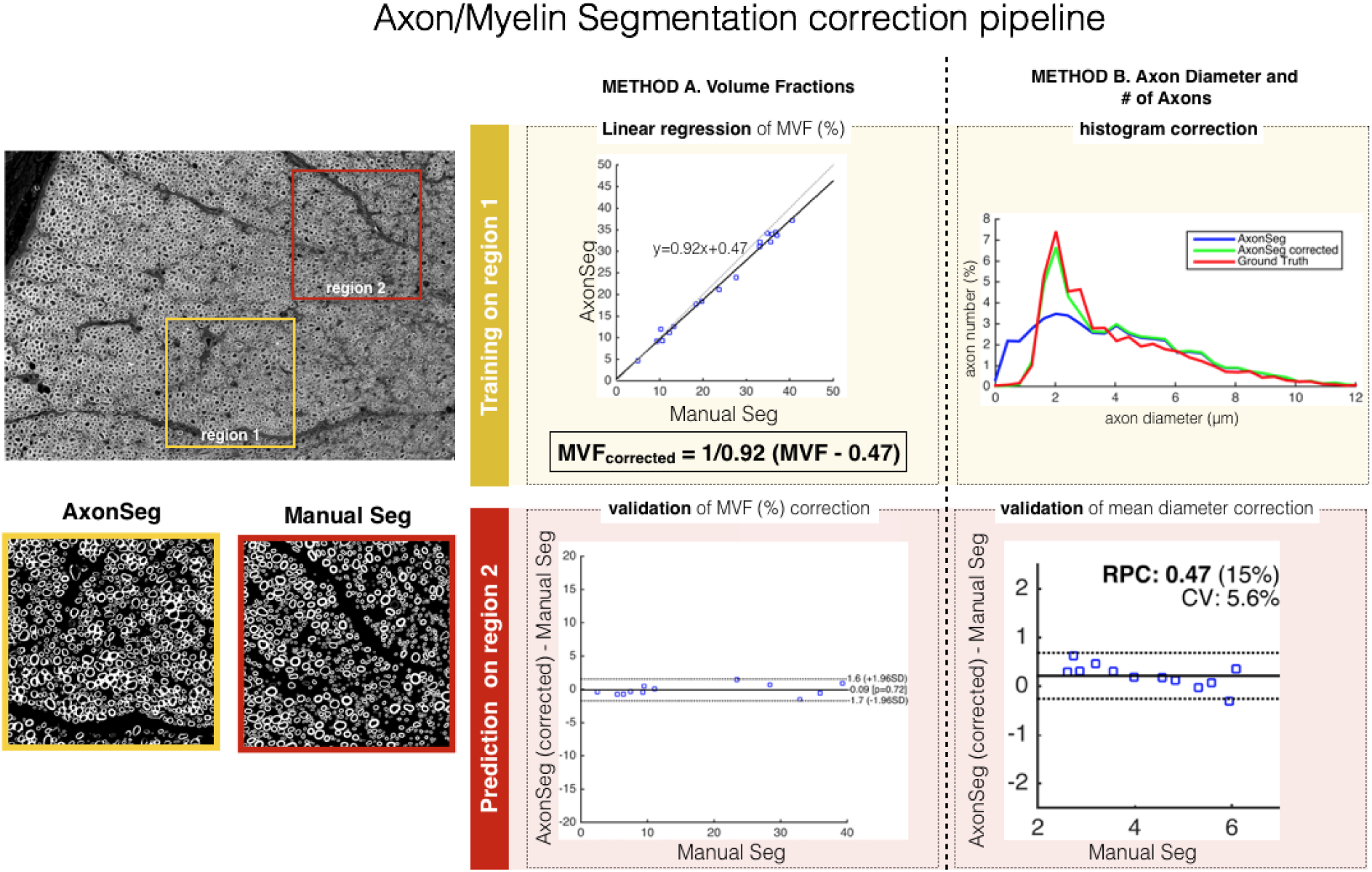
Correction pipeline to compensate for the imperfect segmentation sensitivity and specificity. Top left: Two regions (red and yellow squares) were segmented both manually and automatically with AxonSeg. Bottom Left: Example of myelin segmentation using AxonSeg and Manual segmentation. Top right: The yellow region was used to train the corrections. Bottom right: The red region was used to validate the correction using a Bland-Altman analysis.

As mentioned in the image processing section, the resolution limit was found to be 1µm as the AxonSeg software would mostly segment false positives in that diameter range. Furthermore, from doing this statistical correction, it was observed from the ground truth that there were many axons smaller than 1µm that were not segmented (false negatives).

Figure S6 below shows a comparison of the corrected statistics with the groundtruth using bland-altman plots. This was calculated using a 95% confidence interval (± 1.96*sd). The reproducibility coefficient was 2.2%, 1.6%, 40 and 0.47µm for fiber density, myelin content, number of axons (in 100×100µm^2^ window) and axon diameter respectively.

**Figure S6.**
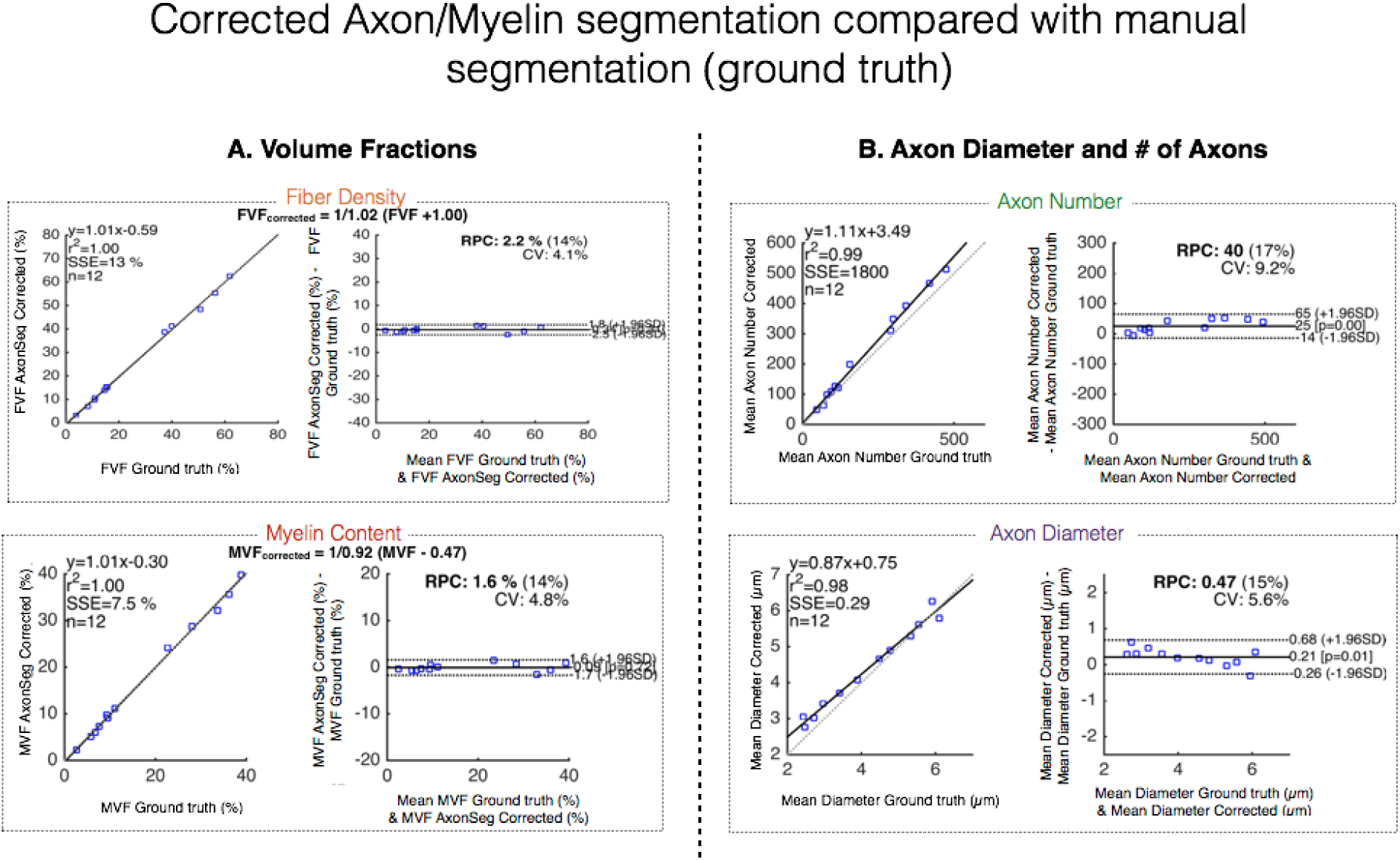
Bland-Altman plots comparing the axon/myelin segmentation (after corrections) with the manual segmentation. The first curve depicts the linear correlation while the second curve shows the agreement between the groundtruth and the segmentation. The RPC is the reproducibility coefficient with the percentage of value next to it. The CV is the coefficient of variation which is essentially the standard deviation of the mean values in percent form.

### Effect of axon orientation

The diameter overestimation for oblique axons can be formulated (assuming that axons are perfect cylinders) as follows. First, the eccentricity *e* of the ellipse obtained by the cutting of the axon is a function of the angle θ of the axon ^48^, and a function of the major (a) and minor (b) semi-axis of the ellipse:

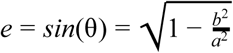

Second, the equivalent diameter *d* (metric reported in this manuscript) of an ellipse is:

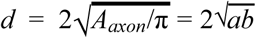

Finally, the equivalent diameter *d* of an oblique axon overestimates the true axon diameter (= 2*b*) by: 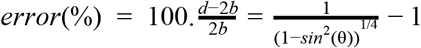. It can be observed in figure S7 that this overestimation exceeds 50% for axons running with an angle of 60° or more. Note that AxonSeg filters highly oblique axons (>73° assuming perfectly cylindrical axon) based on their minor over major axis ratio.

**Figure S7:**
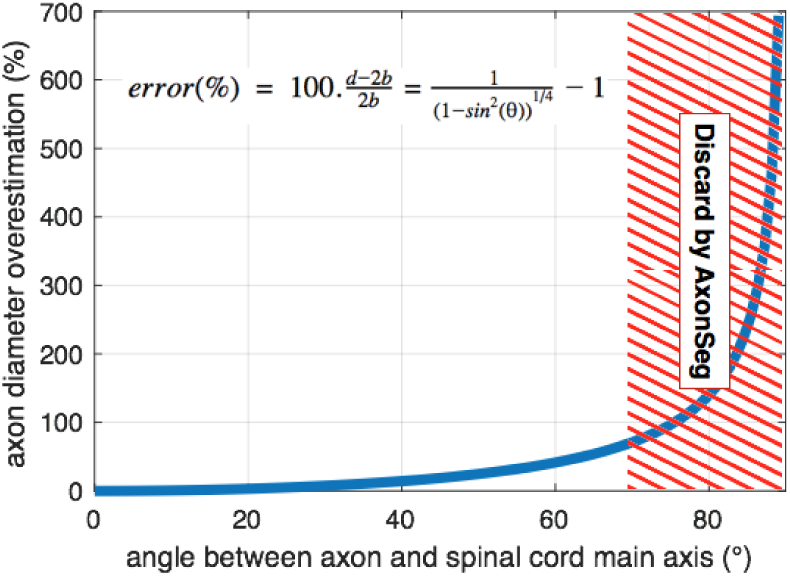
Axon equivalent diameter overestimates the true axon diameter for axons that are not running along the spinal cord main axis. Oblique axons are filtered by AxonSeg based on the minor over major axis ratio of the axon to prevent dramatic failure.

### Segmentation in the gray matter

Although AxonSeg was trained and designed for white matter regions, performance was satisfactory when compared to manual segmentation in the gray matter (see figure S8): in the gray matter, AxonSeg has a sensitivity of 66%, a precision of 64% and <30% error for all metrics except a large overestimation in the number of large (4-8µm) axons (70-80% overestimation). Note that AxonSeg is blind to fibers running perpendicularly to the cord, but these fibers are qualitatively very few.

**Figure S8.**
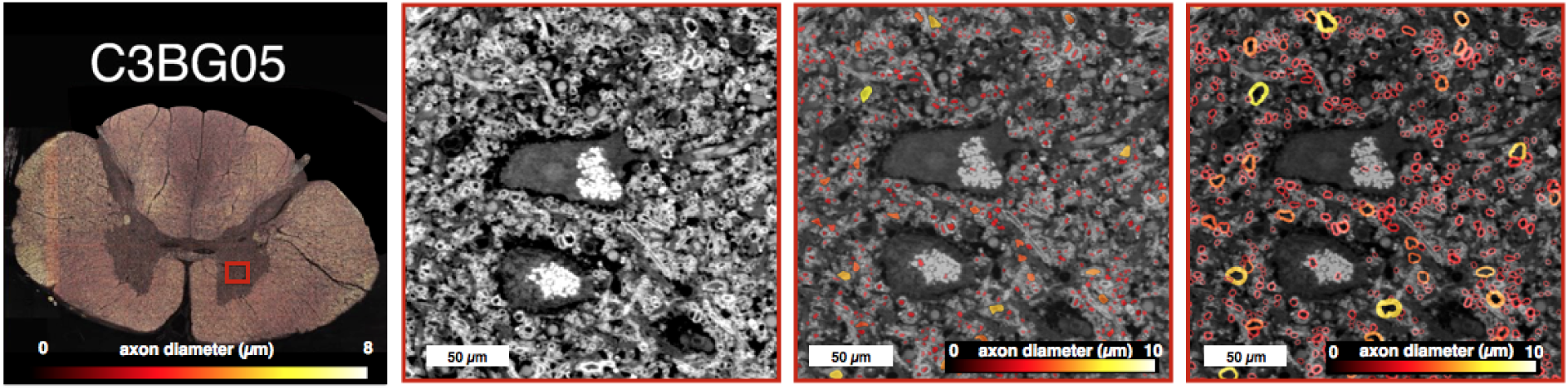
AxonSeg performance in the gray matter. Similar to figure 2 but in a gray matter region. Large axons (4-8µm) and fiber density were overestimated by 70-80%, but other metrics (mean diameter, shape) were quite accurate (<30% bias).

### Laterality difference

Morphometric values extracted per tract in all slices were used to performed a Bland-Altman analysis comparing left versus right tracts (see figure S9). All metrics were found remarkably symmetric with an average difference (± 95% limit of agreement) of 0.01 ± 0.34 µm, 1 ± 9 % and 0.4 ± 6 % for axonal diameter, axonal density and myelin content respectively.

**Figure S9.**
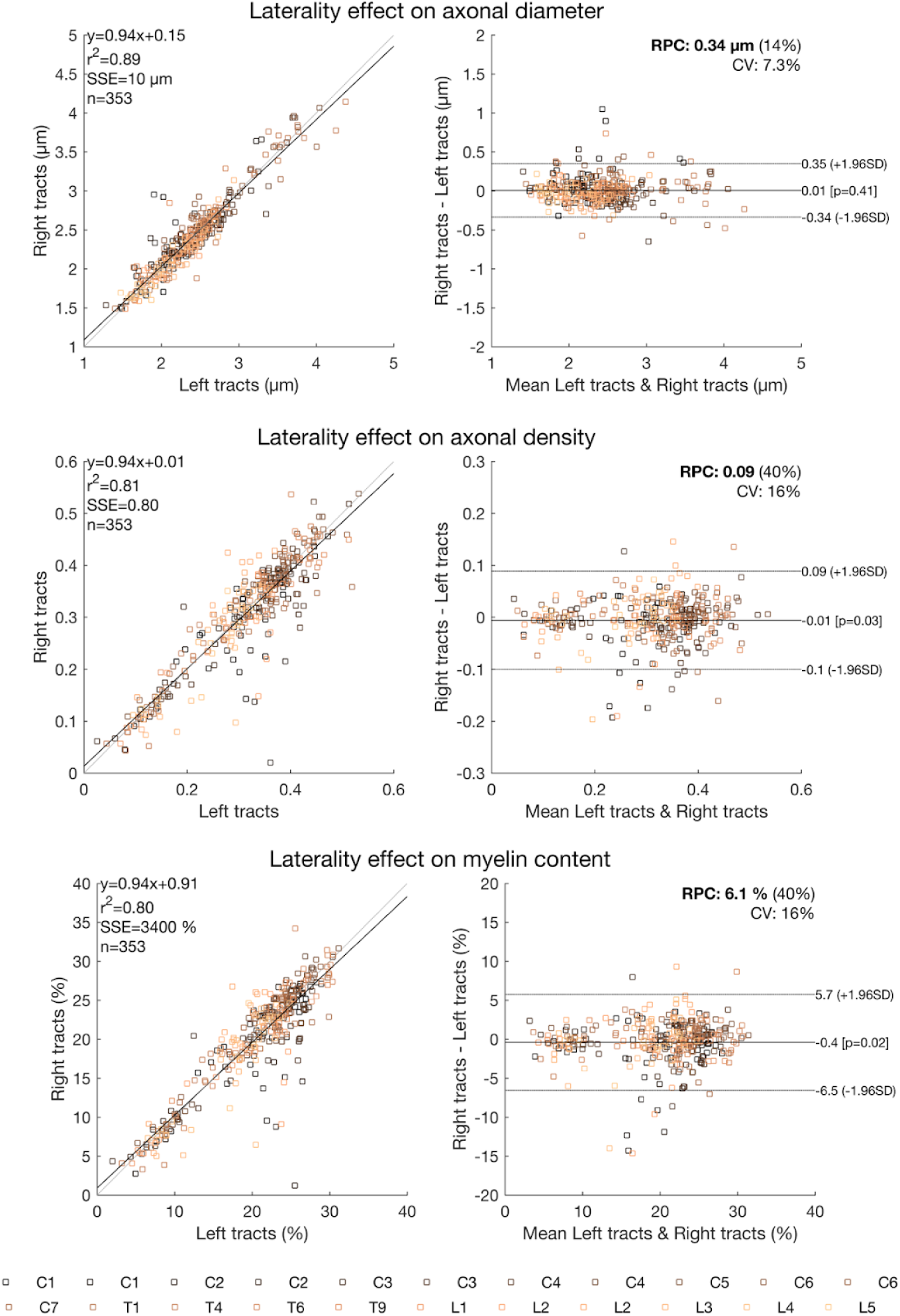
Axon morphometry in the left versus right tracts. Differences of axon morphometry between left and right tracts was very limited: 0.01 µm for axonal diameter, 1% for axonal density, and 0.4% for myelin content.

## Footnotes

1 https://www.mathworks.com/matlabcentral/fileexchange/34943-fit-all-valid-parametric-probability-distributions-to-data?focused=5228686&tab=function

